# Universal Light-Sheet Generation with Field Synthesis

**DOI:** 10.1101/427468

**Authors:** Bo-Jui Chang, Mark Kittisopikul, Kevin M. Dean, Philippe Roudot, Erik Welf, Reto Fiolka

## Abstract

We introduce Field Synthesis, a theorem that can be used to synthesize any scanned or dithered light-sheet, including those used in lattice light-sheet microscopy (LLSM), from an incoherent superposition of one-dimensional intensity distributions. This user-friendly and modular approach offers a drastically simplified optical design, higher light-throughput, simultaneous multicolor illumination, and a 100% spatial duty cycle, thereby providing uncompromised biological imaging with decreased rates of photobleaching.

## Main Text

For years, confocal microscopy was the method of choice for subcellular volumetric imaging of biological specimens.^1^ However, the exceedingly small dwell-times of modern confocal microscopes (e.g., < 2 microseconds) necessitates high peak illumination intensities, which in addition to the excitation wavelength, accelerates photobleaching and phototoxicity.^2, 3^ Further, regions above and below the laser focus are unnecessarily illuminated, resulting in the expenditure of precious photons that do not productively contribute to image formation. In contrast, light-sheet fluorescence microscopy (LSFM) adopts an orthogonal imaging geometry whereby the sample is illuminated from the side and fluorescence is collected at 90 degrees with high numerical aperture optics and million-fold camera-based parallelization.^4^ Because no fluorescence is generated above and below the depth-of-focus, optical sectioning is automatically achieved, and unnecessary illumination of the specimen is avoided. Consequently, LSFM permits unparalleled long-term imaging of dynamic biological specimens.

In LSFM, a critical tradeoff between axial resolution (influenced by the thickness of the light-sheet), field of view (the length over which a beam can approximate a sheet), illumination duty-cycle, and illumination confinement exists. In an effort to simultaneously optimize these optical parameters, the Betzig laboratory developed lattice light-sheet microscopy (LLSM).^5^ Here, the specimen is coherently illuminated with multiple plane waves to sculpt a propagation-invariant 2-dimensional optical lattice. By dithering the lattice (*i.e*., scanning it over a small lateral distance) a time-averaged sheet of light is established. Unfortunately, LLSM requires an optical train that is prohibitively complex, expensive and lossy (~0.1-1% input laser power delivered to the sample).^5^

Here we show that any scanned (or dithered) light-sheet, including lattice light-sheets, can be synthesized by a simple and universal optical process described by a mathematical theorem that we refer to as Field Synthesis (Supplementary Note 1). Applied to Bessel beam light-sheets, Field Synthesis achieves 100% spatial duty cycle, it is more sensitive and results in less photobleaching than time-averaged light-sheets generated by lateral beam scanning. Applied to LLSM, light-losses are reduced 15-fold (Supplementary Note 2) and simultaneous multi-color imaging becomes possible. Furthermore, the number of optical components is decreased 2-fold. Importantly, no spatial light-modulator, diffraction optics or polarization optics are needed, which significantly reduces the complexity and cost of a LLSM system (see Figure S1 and Note 2). We mathematically prove the Field Synthesis theorem, demonstrate proof-of-concept imaging, and provide software and documentation so that the biological community may adopt the technology.

The essence of the Field Synthesis theorem can be described as follows: The projection of the squared modulus of a two-dimensional function is equivalent to the average of the squared modulus of a line scan in the Fourier domain of the same function (Supplementary Notes 1 and 3, Movie S1). Here, the line scan follows the same direction in Fourier space as the projection in real space (*e.g*., k_x_ and x, respectively). Importantly, this has practical applications to scanned light-sheets, where a time-averaged light-sheet is obtained by laterally scanning the illumination beam. In a conventional LSFM, the illumination beam is shaped by light distribution with spatially varying amplitude and phase in the back-pupil plane of the excitation objective, which can be described as 2-dimensional pupil function (Fig 1A, Top Row).^6^ Our theorem predicts that if a focused line is scanned over the same pupil function, an identical time-averaged light-sheet is obtained (Figure 1A, Bottom Row). Thus, by simply scanning a focused line over an appropriate pupil mask (e.g., a 2-dimensional spatial filter), we can create any type of scanned or dithered light-sheet, including those used by LLSM.

**Figure 1.**
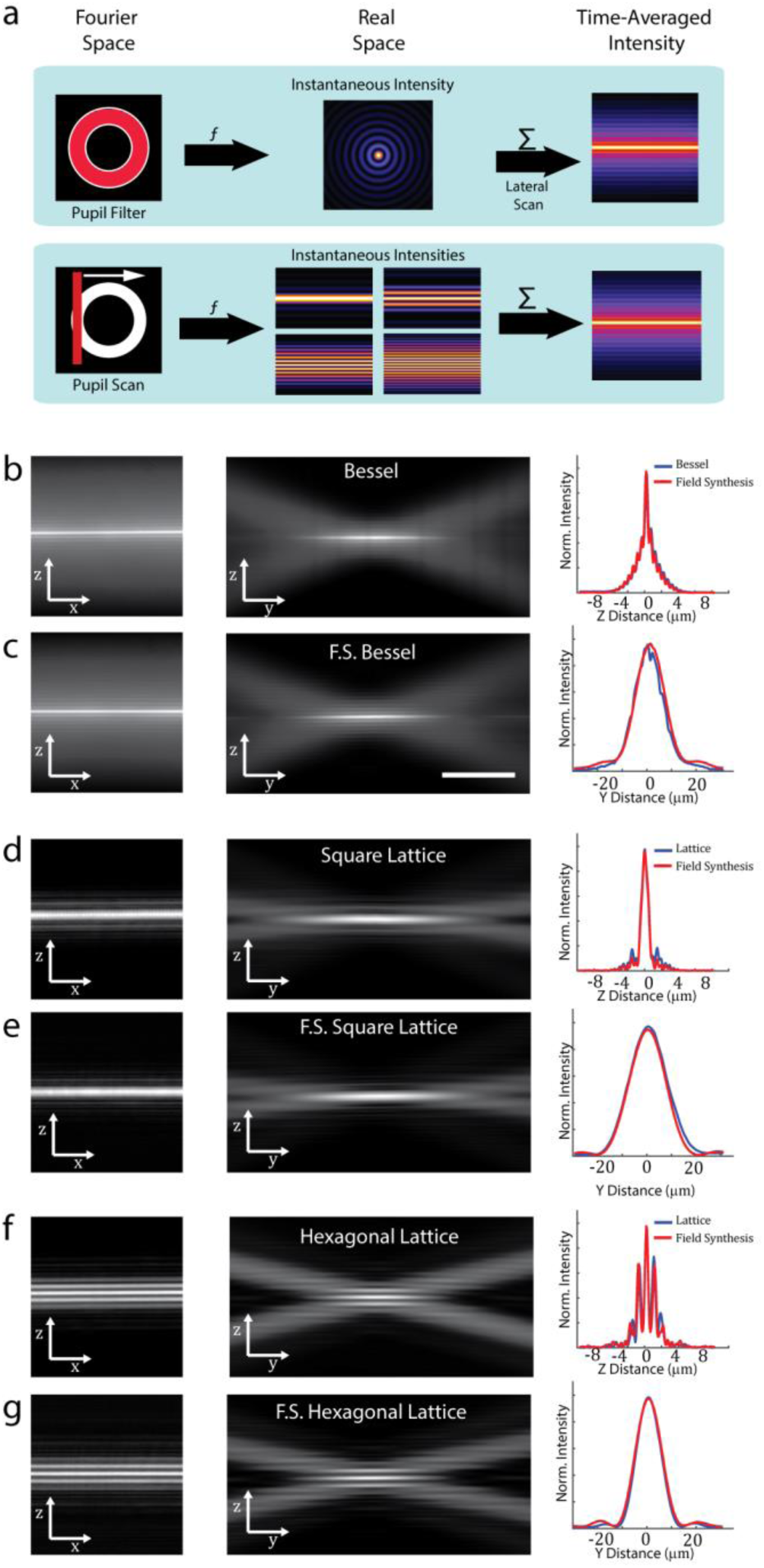
Light-sheet generation by field synthesis. **A, Top**. In light-sheet microscopy, a pupil filter conjugate to the back pupil plane of the illumination objective is used to shape the instantaneous laser focus at the front focal plane of the same objective. To generate a time-averaged sheet of light, the laser focus is rapidly scanned laterally. **A, Bottom**. Principle of field synthesis: a focused line is laterally scanned over a pupil filter, creating a time-averaged sheet of light in the front focal plane of the illumination objective. **B, D, and F**. Experimental examples of a Bessel beam light-sheet, square lattice, and hexagonal lattice, generated by traditional methods, respectively. **C, E, and G**. Experimental examples of a Bessel beam light-sheet, square lattice, and hexagonal lattice generated with Field Synthesis, respectively.

While the time-averaged intensity distributions obtained by Field Synthesis are expected to be identical to dithered or scanned light-sheets, there are important differences: (i) The spatial duty cycle is 100% for symmetric pupil functions (as used in Bessel and LLSM) since the light-sheet is synthesized from a series of cosines (which have maxima at their origin). (ii) Field Synthesis, as implemented here, is based on refraction and hence can be implemented with achromatic optics, permitting simultaneous multi-color excitation.

To evaluate this optical principle, we compared the three-dimensional intensity distributions of Bessel and LLSM with those generated by Field Synthesis (see Supplementary Information for details). As seen in Figures 1B-G, both in cross-sectional views as well as in the propagation direction, field synthesis produces identical light-sheets. Interestingly, for LLSM, it has been hypothesized that complex 2D interference patterns are necessary for its advantageous properties, such as aspect ratio of the sheet and suppression of its sidelobes. Remarkably, we found that the same light-sheets can be obtained by incoherently summing 1D intensity patterns (see Supplementary Notes 1 and 3, Figures S4-S9, Supplemental MATLAB code).

To compare the performance between a Bessel beam LSFM constructed using conventional beam scanning and Field Synthesis, we measured photobleaching curves of 100 nm fluorescent microspheres and microtubule +Plus-End Tracking Proteins (TIPs, labeled with EB3-mNeonGreen) in U2OS cells. In both cases, owing to its higher duty cycle, Field Synthesis resulted in lower rates of photobleaching (Fig. 2 A-C, Movie S2). The low spatial duty cycle of scanned Bessel beams has been previously identified to be the cause for increased photo-bleaching and spurred the development of LLSM.^5, 7, 8^ Our results show a new, general mechanism to improve spatial duty cycle for propagation invariant beams without requiring complex 2D interference patterns and retaining the original light-sheet.^9-12^

**Figure 2.**
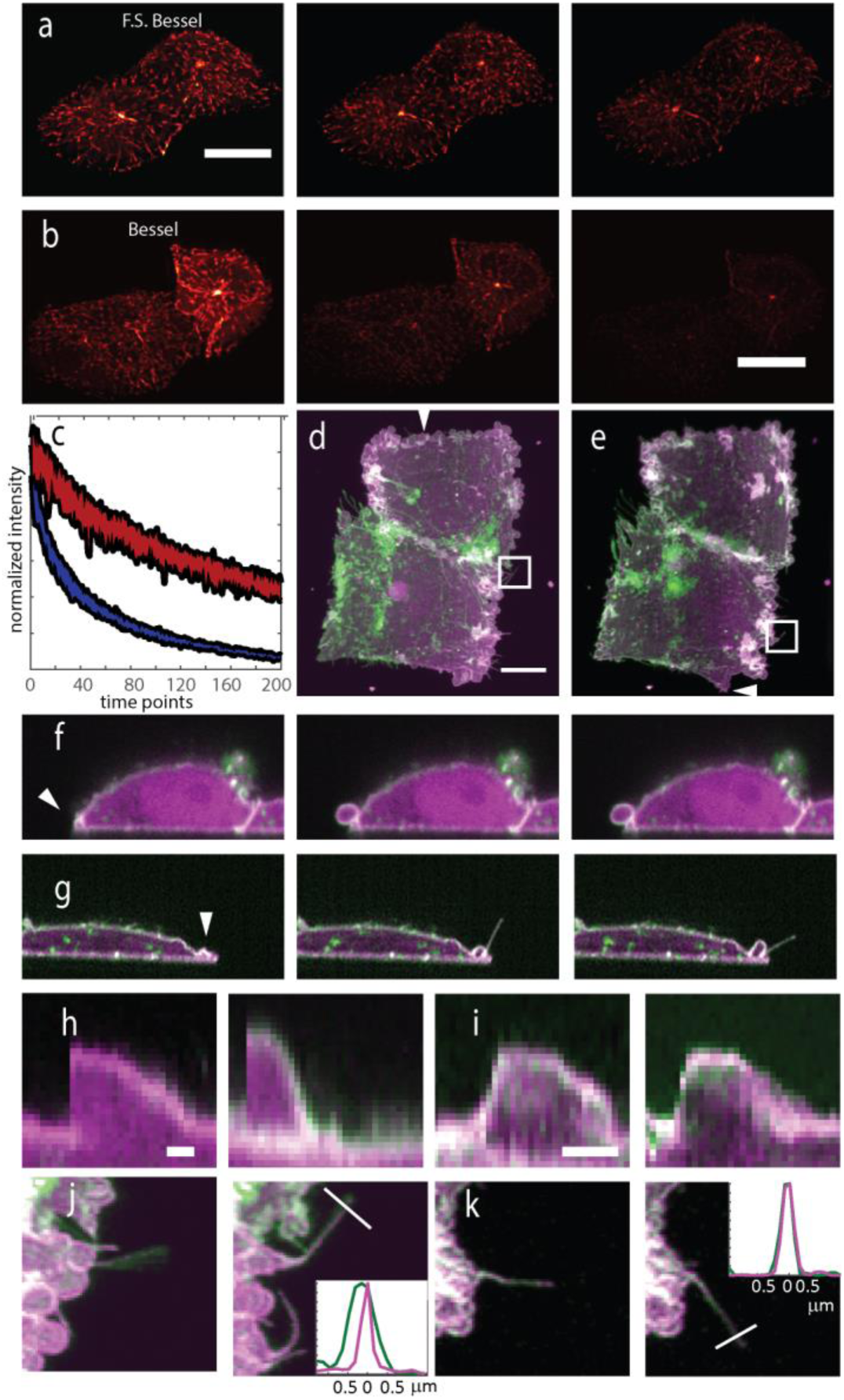
Field Synthesis reduces photobleaching and enables simultaneous multicolor imaging. **A**. Three timepoints (first, fifteenth, and thirtieth) of a movie of EB3 in an U2OS cell as imaged with Field Synthesis. **B**. Three time points (first, fifteenth, thirtieth) of a movie of EB3 in an U2OS cell as imaged with scanned Bessel Beam light-sheet microscopy. **C**. Comparison of photobleaching rate of 100nm fluorescent microspheres over 200 acquired volumes as imaged with Field Synthesis and Bessel Light-sheet microscopy. Shaded bands represent the 95th percentile of each photobleaching curve (11 beads were measured for each condition). **D** MV3 cancer cell as imaged with sequential dual color square lattice light-sheet microscopy at a volumetric rate of 0.23 Hz. Membrane is shown in green and an AKT-PH biosensor is shown in magenta. **E**. Same cell imaged with Field Synthesis in square lattice mode at a volumetric rate of 0.47Hz. **F**. Three consecutive timepoints of a cross-section at position indicated by arrow in **D**. **G**. Three consecutive timepoints of a cross-section indicated by an arrow in **E**. **H**. Kymographs of two blebs imaged with square lattice light-sheet microscopy. Onset of bleb formation is discontinuous. **I**. Kymographs of two blebs imaged with Field Synthesis. Growth of bleb is gradual due to increased sampling frequency. **J**. Magnified view of the boxed region in **D** showing two timepoints of filopodia dynamics. **K**. Magnified view of the boxed region in **E** showing two timepoints of filopodia dynamics. Scale bar: **A**,**B,D**: 10 microns; **H,I**: 20s

To demonstrate that Field Synthesis can be used as an alternative to LLSM, we performed sensitive dual-color imaging of MV3 melanoma cells labeled with a PI3K biosensor (GFP-AktPH) and a membrane-localized fluorescent protein (Td-tomato-CAAX). Both conventional LLSM and Field Synthesis LLSM allowed us to resolve fine cellular features such as small membrane protrusions and filopodia and provided excellent optical sectioning. However, because Field Synthesis is compatible with simultaneous multicolor imaging, the volumetric imaging rate is two-fold faster than conventional LLSM if two-channels are acquired, despite using identical imaging parameters (camera integration time, z-step size, etc.). This allowed us to observe rapid subcellular events such as small membrane protrusions and filopodia dynamics, which were temporally undersampled in the sequential acquisition mode afforded by conventional LLSM (Figure 2d-k, Movie S3-S5). While sequential LLSM acquisition could in principle be accelerated by using shorter camera integration times, this would come at the expense of loss of signal or higher peak illumination intensities and might not be possible due to camera bandwidth limitations. Thus we believe that simultaneous multicolor imaging with Field Synthesis expands the capabilities of LLSM, particularly in spectral imaging applications.^13^

In conclusion, we have introduced with the Field Synthesis theorem a novel and universal mechanism to generate scanned and dithered light-sheets. Importantly, Field Synthesis drastically simplifies the optical train compared to lattice light-sheet microscopy. The consequence is that there are less critical components to align and temporally control, which eases building and operating such an instrument, both practically and financially. Additionally, simultaneous multicolor acquisition can be achieved. Beyond LLSM, the field synthesis principle applies to any type of intensity distribution. So, by simply changing the pupil filter, other light-sheet types can be generated, including, Bessel, Gaussian, or Airy light-sheets. Our insight that any light-sheet can be synthesized by a superposition of one-dimensional functions also implies that field synthesis could be used to rationally design new forms of light-sheets, each uniquely optimized to specific merit functions. Thus, we believe that Field Synthesis will make light-sheet fluorescence microscopy modular and democratize advanced techniques, including but not limited to lattice light-sheet microscopy.

## Acknowledgements

We would like to thank the Cancer Prevention Research Institute of Texas (RR160057 to R.F.), the National Institutes of Health (T32CA080621 to M.K., F32GM117793 to K.M.D., K25CA204526 to E.S.W.), and Human Frontier Science Program (LT000954/2015 to P.R.).

## Author Contributions

R.F., K.M.D., and B-J.C. designed the research. B-J.C. and R.F. designed, built and operated the microscope. R.F., K.M.D., P.R., and B-J.C. performed image analysis. E.W. provided the MV3 cancer cells. M.K. carried out mathematical derivations, proved the theorem, and revised the MATLAB code. R.F., K.M.D., and B.-J.C wrote the manuscript, and all authors read and provided feedback on the final manuscript.

## Additional Information

The datasets acquired for this study are available from the corresponding author upon request.

## Supplementary Information

### Microscope Setup

The basic units to generate lattice and field synthesis light-sheets are shown in Figure S1 and our microscope setup is illustrated in Figure S2. A 488 (Sapphire 488–300 LP, Coherent) and a 561 nm (OBIS 561–150LS, Coherent) are combined with a dichroic mirror (LM01–503–25, Semrock), and focused through a 25-micron spatial filter (P25H, ThorLabs) with a 50 mm focal length achromatic lens (AC254–050-A, Thorlabs). Thereafter, flip mirror 1 is used to select either the LLSM or Field Synthesis optical paths, and the beam is recollimated with a 50 (AC254–050-A, Thorlabs) or 400 (AC508–400-A, Thorlabs) mm focal length achromatic lens, respectively. Further, the optical system is operated in a fluorescence (e.g., with orthogonal detection, Figure S2A) or transmission (e.g., with diascopic detection, Figure S2B) mode, to image fluorescent specimens and investigate light-sheet properties, respectively.

The LLSM path is a simplified version of the original set-up.^1^ The collimated and expanded beam is further magnified and shaped to a line beam by a pair of achromatic cylindrical lenses (68–160, Edmund optics and ACY254–200-A, Thorlabs). The line beam uniformly illuminates a region on the spatial light modulator (SLM, SXGA-3DM, Forth Dimension Displays) where the lattice pattern is displayed. A polarized beamsplitter (10FC16PB.3, Newport) and a half-wave plate (AHWP10M-600, Thorlabs) are placed in front of the SLM, and together with the SLM form the pattern generator^12^. The lasers were assembled in a way that the polarization states fit the pattern generator. The diffraction light from the SLM is focused by an achromatic lens (AC508–400-A, Thorlabs) on a custom-designed mask (mask 1, photo sciences). The mask has various annuli to block unwanted diffraction orders (mainly 0 order but also higher orders). Different sizes of the annuli correspond to different outer and inner NAs on the back focal plane of the illumination objective lens. After passing through the mask, the desired diffraction orders are de-magnified through two achromatic lenses (AC254–150-A, AC254–100-A, Thorlabs) and fit into the size of a galvanometer (Galvo mirror, 6215H, Cambridge Technology). A folding mirror, an achromatic lens (AC-254–100-A, Thorlabs), the flip mirror 2 and a tube lens (ITL-200, Thorlabs) then direct the light to the illumination objective lens (40X, NA0.8, 3.5mm WD, CFI Apo 40XW NIR, Nikon) to form a light-sheet in sample space. All distances between a pair of lenses equal to the summation of their focal lengths so any pair of lenses forms a 4f arrangement. Thus, The SLM is conjugated to the focal plane of the illumination objective lens. The mask is at the Fourier plane of the SLM and is conjugated to the back focal plane of the objective lens. As the galvanometer is conjugated to the mask and the back focal plane of the objective lens, it scans the light-sheet laterally (y) on the focal plane of the detection objective, i.e. enables rapid lateral scanning of a Gaussian or Bessel light-sheet and the dithered mode of the lattice light-sheet.

The Field synthesis path consists of an iris (IDA20, Thorlabs), an achromatic cylindrical lens (ACY254–050-A, Thorlabs), an achromatic lens (AC254–075-A, Thorlabs), another custom-designed mask (mask 2), and a tube lens (ITL-200, Thorlabs). Additionally, a galvanometer (Galvo mirror, 6215H, Cambridge Technology) and a folding mirror are placed in between the achromatic cylindrical lens and the achromatic lens. This path shares the same final tube lens and illumination objective lens with the lattice LSFM path. All distances between a pair of lenses equal to the summation of their focal lengths so any pair of lenses forms a 4f arrangement. The galvanometer in the Field Synthesis path is conjugated to the focal plane of the illumination objective lens. The mask is conjugated to the back focal plane of the objective lens, which is conjugated to the same plane as the mask in the Lattice LSFM path. Since the magnification between the mask and the back focal plane of the objective lens is different in the Field Synthesis LSFM path and the Lattice LSFM path, the sizes of the annuli on mask 2 are designed to match the outer and inner NAs resulted from the annuli on mask 1, as shown in Table 1.

The detection path is orthogonal to the illumination path as in a conventional LSFM set-up and it consists of an detection objective lens (40X, NA0.8, 3.5mm WD, CFI Apo 40XW NIR, Nikon), a tube lens (ITL-200, Thorlabs), a dichroic mirror (Di03-R561-t3–25⨉36, Semrock) that separates the green and the red fluorescence, and two identical sCMOS cameras (Orca Flash4.0 v2, Hamamatsu). A red fluorescence emission filter (FF01–593/LP-25, Semrock) and a green fluorescence emission filter (ET525/50m, Chroma) is placed in front of camera 1 and camera 2, respectively. A piezo stage (P621.1CD, Physik Instrumente) is used to move the sample diagonally to the detection axis in the fluorescence mode (fig. S2A) or move the detection objective lens in the transmission mode (fig. S2B).

In the transmission mode (fig. S2B), the detection objective lens is placed along the propagation of the light-sheet and an achromatic lens (AC508–400-A, Thorlabs) is used as the tube lens. An ND filter (ND40A, Thorlabs) is placed in front of the sCMOS camera (Orca Flash4.0 v2, Hamamatsu) to reduce the intensity of the straight light from the lasers. The combination of the 400mm tube lens and the 40x objective lens offers a final magnification of 80x, which results in one pixel on the camera corresponds to 81.25 nm and that can present the light-sheet with a reasonable contrast and resolution. By stepping the detection objective lens with the piezo stage through the illumination lattice, we investigate and measure the 3D light-sheet properties.

The pattern on the back focal plane of the excitation objective lens can be observed by Camera 4 (DCC1545M, Thorlabs) with an ND filter (ND10A or ND20A, Thorlabs) in the front when flipped mirror 3 is used. Two achromatic lenses (AC254–100-A and AC254–050-A, Thorlabs) are used to image the back focal plane of the objective lens, which is conjugated to the plane of the masks, on the camera. By observing the pattern on this plane allows us to verify the final outer and inner NAs from both masks and adjust the scan range of the galvo in Field Synthesis.

### Microscope control and image acquisition

The data acquisition computer was equipped with an Intel Xeon E5–2687W v3 processor operating at 3.1GHz with 10 cores and 20 threads, 128GB of 2133MHz DDR4 RAM, and an integrated Intel AHCI chipset controlling 4x 512GB SSDs in a RAID0 configuration. All software was developed using a 64-bit version of LabView 2016 equipped with the LabView Run-Time Engine, Vision Development Module, Vision Run-Time Module and all appropriate device drivers, including NI-RIO Drivers (National Instruments). Software communicated with the camera (Flash 4.0, Hamamatsu) via the DCAM-API for the Active Silicon Firebird frame-grabber and delivered a series of deterministic TTL triggers with a field programmable gate array (PCIe 7852R, National Instruments). These triggers included analog outputs for control of mirror galvanometers, piezoelectric actuators, laser modulation and blanking, camera fire and external trigger, and triggering of the ferroelectric spatial light modulator. A timing diagram for the galvanometric beams scanning in Field Synthesis illustrated in Fig. S3. All images were saved in the OME-TIFF format.

Some of the core functions and routines in the microscope control software are licensed under an MTA from Howard Hughes Medical Institute, Janelia Farm Research Campus. The control software code for Field Synthesis can be requested from the corresponding authors and will be distributed under MTA with HHMI Janelia Research Campus.

### Data Analysis

Image analysis was performed with Fiji^3^ and MATLAB (Mathworks). To evaluate the rate of photobleaching for 100 nm fluorescent microspheres, the average intensity of 11 similarly bright beads were measured through time by recording their peak intensity at every timepoint. To this end, a Matlab script was written that detected the peak intensity of all beads in the dataset and a subset was manually selected that yielded the same average initial intensity for both imaging modalities. The same script was used to initially calibrate the power levels that yielded similar fluorescent counts when the same sample region was imaged with either Bessel beam or Field Synthesis light-sheets. Both the bead imaging as well as the EB3 imaging in the U2OS cells was conducted with such calibrated power levels.

### Deconvolution and data post processing

The point spread function (PSF) used for deconvolution of the lattice light-sheet (conventional and Filed Synthesis) data was synthesized by using the PSF Generator plugin in ImageJ.^3,4^ The Born and Wolf 3D optical model is used to generating the PSF. Illumination and detection PSFs are synthesized in 3D with the practical experimental parameters (i.e. the refractive index, the wavelength, the NA, and the voxel size). The final light-sheet PSF is the product of the illumination and detection PSFs. Since our sample is placed diagonally to the illumination and detection and we scan the sample to acquire the images (sample-scan acquisition), the light-sheet PSF is sheared oppositely referring to the direction of the sample scan. 3D-deconvolution was performed with 10 iterations of the Richardson-Lucy algorithm as implemented in MATLAB. The deconvolved data was sheared at an angle of 45 degrees using an affine transform and bleach corrected with Fiji using the histogram matching method.

For microtubule +TIP data, experimental point-spread functions were obtained by imaging isolated 100 nm nanospheres with both the conventional Bessel and Field Synthesis modes of microscope operation. We chose experimental PSFs here as the Bessel beam light-sheet has a more complex structure and hence could not be approximated with the numerical PSF Generator. These data were deconvolved, the Z-axis bicubically interpolated so that the voxel dimensions were isotropic (162.5 nm) and sheared at an angle of 45 degrees using an affine transform. The EB3 data was not bleach corrected.

### Biological Sample Preparation

Both metastatic melanoma (MV3) and osteosarcoma (U2OS) cell lines were cultured in DMEM supplemented with 10% fetal bovine serum and penicillin/streptomycin and maintained at 37 degrees Celsius with 5% CO^2^ atmosphere. MV3 cells were lentivirally transduced using the pLVX-IRES-PURO and pLVX-IRES-NEO expression systems. U2OS cells were lentivirally transduced with a modified pLVX-shRNA2 vector with a truncated CMV promoter to reduce ectopic expression, as previously described.^5^ All cells were plated on 5 mm coverslips and mounted in a custom sample holder for imaging.^6^

### Quantification of Beam Geometry

For direct measurements of the light-sheet intensity distribution, the excitation light was imaged in transmission with a 40x water-dipping objective, which was scanned in the Y-direction (e.g., along the optical axis) with a 100-micron piezo actuator. Using a 400 mm tube lens (Fig. S2B), the effective pixel size of the imaged beam was 81.25 nm. Beam analysis software, written in MATLAB, identified the laser focus in Y, the size of the beam waist in Z and the beam propagation length in Y. Since Bessel and lattice light-sheets are propagation invariant (i.e. they maintain their beam profile over extended propagation distances), we did not apply the Rayleigh length criterion to characterize the light-sheets. Instead, we use the full width half maximum in the propagation direction.

### Sequential Lattice data

We developed a simple experiment, which can be reproduced by anyone who has access to a lattice light-sheet microscope, to test the Field synthesis theorem. We termed the procedure sequential lattice: Here, we used the lattice light-sheet illumination path in the square lattice mode and manually positioned a mask to only transmit isolated diffraction orders along the kz direction (see also fig. S4). This was repeated for three mask positions and the corresponding datasets were added numerically. As the summing of the intensity is incoherent, at no point a 2D lattice pattern emerges, but only one-dimensional intensity patterns are created. Compared to the dithered square lattice, the sequential lattice approach produces the same light-sheets (see fig S4). In particular, some sidelobe structures are more closely reproduced than in Figure 1 d & e. We believe that these small differences arise from slightly different intensity distributions in the pupil plane. Overall, this further confirms that a dithered lattice light-sheet can be synthesized by purely one-dimensional functions, as predicted by the Field Synthesis theorem.

### Fluorescent measurement of light-sheet properties

We have characterized the light-sheets using transmission measurements (Fig. 1 and S2B), which provided images of the light-sheets in sample plane. Due to the involved imaging operation, blurring of the observed light-sheet can be expected, but we argue that this effect will equally affect the conventional and Field Synthesis light-sheets. Nevertheless, we also conducted a direct measurement of the light-sheet in sample space. To this end, we mechanically scanned a 50nm fluorescent bead in 0.2micron steps in the Z-direction and in one-micron steps in the Y-direction through a light-sheet. At every scan position, a widefield image of the fluorescent emission from the bead was taken and the signal on the camera was summed up on a local window surrounding the bead. The window size was chosen such that it contained all diffraction rings of the bead at the largest defocus it was undergoing during the scan. The obtained intensity distributions of a square lattice and a Field Synthesis square lattice light-sheet are shown in Figure S6. The width of the light sheet at its waist was 0.85 microns for the square lattice and 0.91 microns for the Field synthesis light-sheet. In the propagation direction, the Full width half maximum for the square lattice light-sheet was 16.7 microns and 17.2 microns for the Field Synthesis light-sheet, respectively. These values are in close agreement with the transmission measurements shown in Table 2 and the blurring in the transmission measurement appears to be small.

## Supplementary Notes

### Supplementary Note 1 - Field Synthesis Theorem and Proof

The Field Synthesis Theorem that we have formulated underlies the operation of the field synthesis microscope and details how a time-averaged illumination pattern equivalent to any scanned or dithered light-sheet can be synthesized by conducting line scans over a mask in Fourier space.

We show that an ideal line scan in the Fourier domain results in a 2-D intensity function that varies only in one dimension and that this variation is determined by a 1-D Fourier transform of the line scan. This is because each line scanned in the Fourier domain results in a projection in the illumination domain. Meanwhile any phasing created by the location of the line scan does not contribute to the intensity.

The time average of the intensity of the individual line scans is shown to be equivalent to a projection of the original field. Due to Parseval’s Theorem applied in 1-D, summation over the square modulus of each frame of the scan is equivalent to a projection of the square modulus of 1-D inverse Fourier transform of the filter in the Fourier domain. Therefore, it is theoretically possible to synthesize a light-sheet, which is the projection of a filter pattern by doing a time average of a line scan of that filter pattern in the Fourier domain.

#### Field Synthesis Theorem

Let 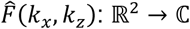 be an arbitrary 2-D amplitude or phase mask at the front focal plane of the lens as in Figure 1A, top. If 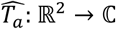, a vertical line scan of this mask at column position *a*, is modeled as the product of 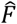 and a 1-D delta function in terms of *k_x_*,

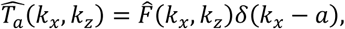
then the sum of the intensity, the square modulus, of the resulting illumination patterns over the columns of the mask is equivalent to the intensity of a lateral scan of the illumination pattern produced by the original mask, within a scalar scaling constant, N,

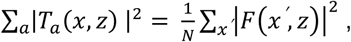
where *T_a_* and *F* are the 2-D inverse Fourier transforms of 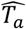 and 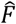, respectively. The prime is included for clarity only.

#### Proof

To determine the form of *T_a_*, the illumination pattern produced by the line scan (Figure 1A and Movie S1, bottom), we want to express it in terms of the illumination pattern, *F*(*x,z*), that results from the instantaneous intensity of the mask (Figure 1A and Movie S1, top).

First, we use substitution to translate the expression of the vertical line scan so that the delta function is expressed as a central slice.

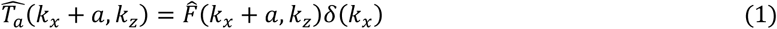

Applying the 2-D Fourier Transform, we use the Fourier shift theorem to express the mask in the illumination domain. This results in the convolution of the modulated illumination pattern and a 1-D delta function in terms of *z*. Note that the complex exponentials modulate the phase but not the amplitude in terms of x. N denotes the number of samples in the x-direction.

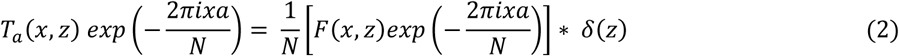

The convolution of illumination pattern with a 1-D delta function in terms of z smears the pattern in the x direction. The smearing results in a summation due to superposition and thus obviates the need for lateral scanning of the pattern.

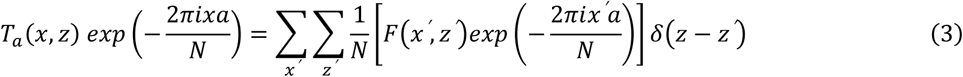

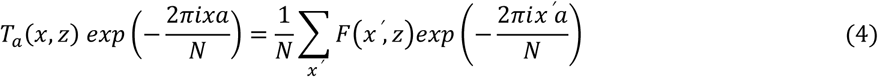

The form of the right-hand side is equivalent to a discrete 1-D Fourier transform of the illumination pattern with respect to x evaluated at *k_x_* = *a*. This is also equivalent to a discrete 1-D inverse Fourier transform of the mask with respect to *k_z_*.

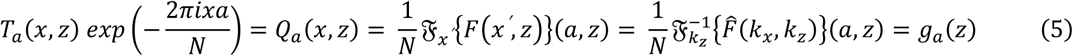

Thus, with the exception of modulation of phase in the x direction on T_a_(x,z), the resulting illumination pattern, Q_a_(x,z), only varies in the z-direction and can be expressed in terms of the 1-D function, 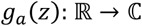. Taking the square modulus to obtain the intensity of the illumination pattern and noting the phase modulation does not affect the amplitude or the intensity, we obtain the following expression for the intensity of the illumination pattern.

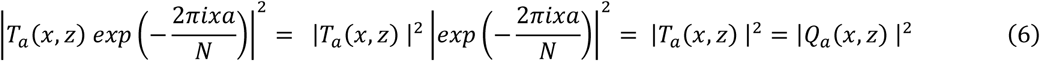

The resulting intensity of the illumination pattern of the vertical line scan of the mask thus only varies in terms of z (Figure 1A and Movie S1, bottom).

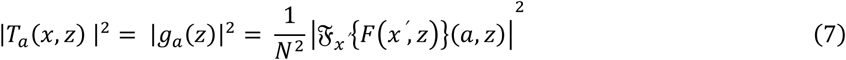

The time averaged intensity pattern is determined by a summation of the intensity of individual line scans over each column *k_x_* = *a*.

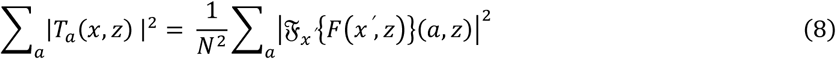

We express the right side of the above equation in terms of the instantaneous intensity of the illumination pattern produced by the original mask, *F*(*x,z*), by applying Parserval’s Theorem in one dimension.

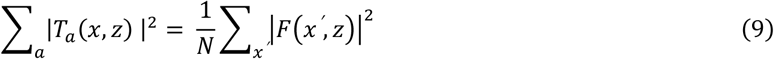

Note that the intensity of the time averaged illumination pattern only varies in the z direction (Figure 1A and Movie S1, bottom). This property is inherited from the individual line scans, which also do not vary in the z direction (Figure 1A and Movie S1, bottom). Also, note that the expression of the right-hand side is equivalent to expression for a lateral scan of the instantaneous intensity of the mask (Figure 1A and Movie S1, top) except for a scalar scaling constant. We have thus proved what we original sought.

#### Discussion of Field Synthesis Theorem Proof

The direct consequence of the proof is that the summation of individual line scans results in the same field pattern that could be achieved by a summation over the original field that could be achieved by scanning or dithering.

In the MATLAB function in *FieldSynthesisTheorem.m* (see Supplemental MATLAB Code), we demonstrate that this is a general result that can be applied to an arbitrary complex mask. The default example for the illumination pattern is the stock cameraman.tif with an added imaginary component. Examining the vertical 1D profile of the generated field with *FieldSynthesisTheorem.m*, we also see that the profile can be calculated either through projection (right side of Equation 9), slicing the pupil mask in the Fourier domain (left side of Equation 9), or applying a 1D dimensional Fourier transform (right side of Equation 8). Taking the vertical profile of the smeared field results is the same operation as the projection method. In the same function, we also show that the square modulus of an individual scan line results in a pattern that only varies in one dimension (Equation 7), but that the electric field is modulated based on the location of the line scan (Equation 5). An interactive version of this is demonstrated in *FieldSynthesisInteractive.m*.

We apply the proof to the comparison between the traditional and field synthesis methods to generate dithered lattice light-sheets in *FieldSynthesisVersusLattice.m*. There we again show the predicted profiles generated by traditional dithering and the field synthesis method are identical. Examining the Fourier transform of the intensity, the autocorrelation of the electric field, we see that while the lattice light sheet pattern has a complex pattern with lateral and diagonal components, the dithered intensity has a very simple and nearly one-dimensional pattern. This is only approximately one-dimensional since there is only limited support for dithering operation and this occurs only over one period of the lattice. This result is suggested by the proof since the resulting pattern only requires a one-dimensional Fourier transform and only varies in the vertical dimension. Additionally, the proof suggests that several masks could result in the same pattern since the only constraint is that the sum of the square modulus of the line scans must equal to the one-dimensional vertical intensity profile.

Overall, the proof suggests that the field synthesis method of constructing a light sheet pattern presents an alternative to scanning or dithering to create a light-sheet. The proof shows that there is considerable flexibility in the choice of a mask to create the same pattern generated by scanning or dithering.

We chose the name Field Synthesis as it was inspired by previous work by Lanni et al^7^ on standing wave microscopy. While our Field Synthesis approach is more general, the generation of a Bessel beam light-sheet resembles the earlier implementation by Lanni et al.

### Supplementary Note 2 - Reduction in light loss and optical complexity

To analyze light-loss and complexity, we focus here only on the optical components that create the lattice and field synthesis light-sheets, which are shown in figure S1. Both units have the same input, a line-shaped intensity distribution, and as an output, generate a lattice light-sheet in an image plane. The layout and components of the lattice light-sheet correspond to the initial lattice microscope publication^1^ and represent to our best knowledge what users will have available (homebuilt or commercial systems).

The fundamental losses in the lattice unit stem from the ferroelectric SLM’s ability to rotate the input polarization and from diffraction losses. Experimentally, we measured 2.4% light throughput for a square lattice pattern through the unit as defined in figure S1. In practice, lower transmission powers have been reported for an entire lattice setup, which involves further light-losses for beam expansion, power modulation and transmission losses through additional optics.

For Field Synthesis, the losses are governed by the transmission through the pupil mask, no SLM and diffraction losses apply. When we adjusted the input beam size with an iris (see also figure S2) so that it matched size of the ring mask, we measured a transmission of 35.4% in the square lattice mode. Thus, for the fundamental light losses between the two techniques, we estimate a ~15 fold higher transmission efficiency for the Field Synthesis method.

When we measured the overall transmission from laser source to the sample, the field synthesis setup was 8.5-fold more efficient then the lattice light-sheet setup. The difference to the fundamental losses stems mainly from the fact that we over-expanded the beam for Field Synthesis to yield a very uniform illumination of the sample (the transversal illumination profile for lattice is more Gaussian in shape). It is possible to tune the transmission properties for both lattice and Field synthesis by modifying the beam expansion. However, the fundamental losses outlined above will put a hard limit on what can be achieved.

Counting the number of optical components in the lattice light-sheet unit in figure S1, there are 10 parts (polarizing beam splitter, waveplate, spatial light modulator, galvo mirror, folding mirror, pupil mask, four lenses), two of which are optical modulators/actuators. In the field synthesis unit, there are five optical components (galvo, folding mirror, lens mask, lens), of which one is an optical actuator.

### Supplementary Note 3 - Discussion on dithering lattice light-sheets

Line scanning of the mask prevents components of the mask from interfering with each other horizontally or diagonally – a combination of horizontal and vertical interference patterns. However, we note that the dithering operation in the formation of convention lattice light-sheets eliminates any contribution of horizontal interference patterns. The dithering operation results in a projection of the illumination pattern and a replication of that projection across the field.

We would like to understand the relationship between the power spectrum of the original illumination field and its dithered projection. By dithering, we average together any intensity variation in the x-direction by scanning the beam across the light-sheet. Since there is no longer any intensity variation in the x-direction and the intensity is now equal to the mean, the illumination intensity of each z-plane is described by a single value, its mean. We thus expect the power spectrum to be a 1D vector at *k_x_* = 0 that only describes the variation along the z-axis.

Formally, the power spectrum of a field *F*(*x,z*) is the Fourier Transform of the intensity field |*F*(*x,z*)|^2^. The power spectrum is evaluated as follows.

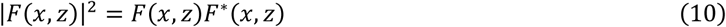

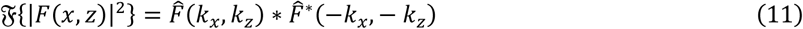

The effect of dithering in the x-direction is equivalent to taking a projection and replicating it across one period of the lattice, P. This in turn can be expressed as convolution with a 1D rect function of width P windowed by a 1D delta function in z.

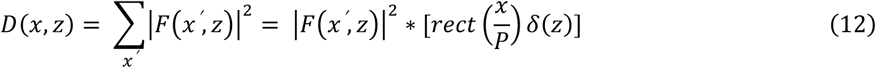

The Fourier Transform of the dithered signal can be then expressed by using the convolution theorem and noting that the 2D Fourier Transform of 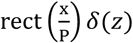 is 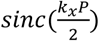.

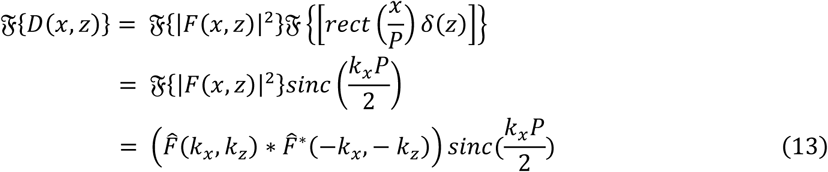

The *sinc* function effectively permits a thin line around *k_x_* = 0. Note that 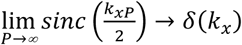 meaning that dithering over a long period will make the window approach the thinnest possible line, *δ*(*k_x_*).

Thus, by dithering the illumination pattern, we select for only the components near *k_x_* = 0 of the power spectrum of the field.

### Supplemental MATLAB Code is available at

https://github.com/AdvancedImagingUTSW/FieldSynthesis

## Supplementary Figures

**Figure S1.**
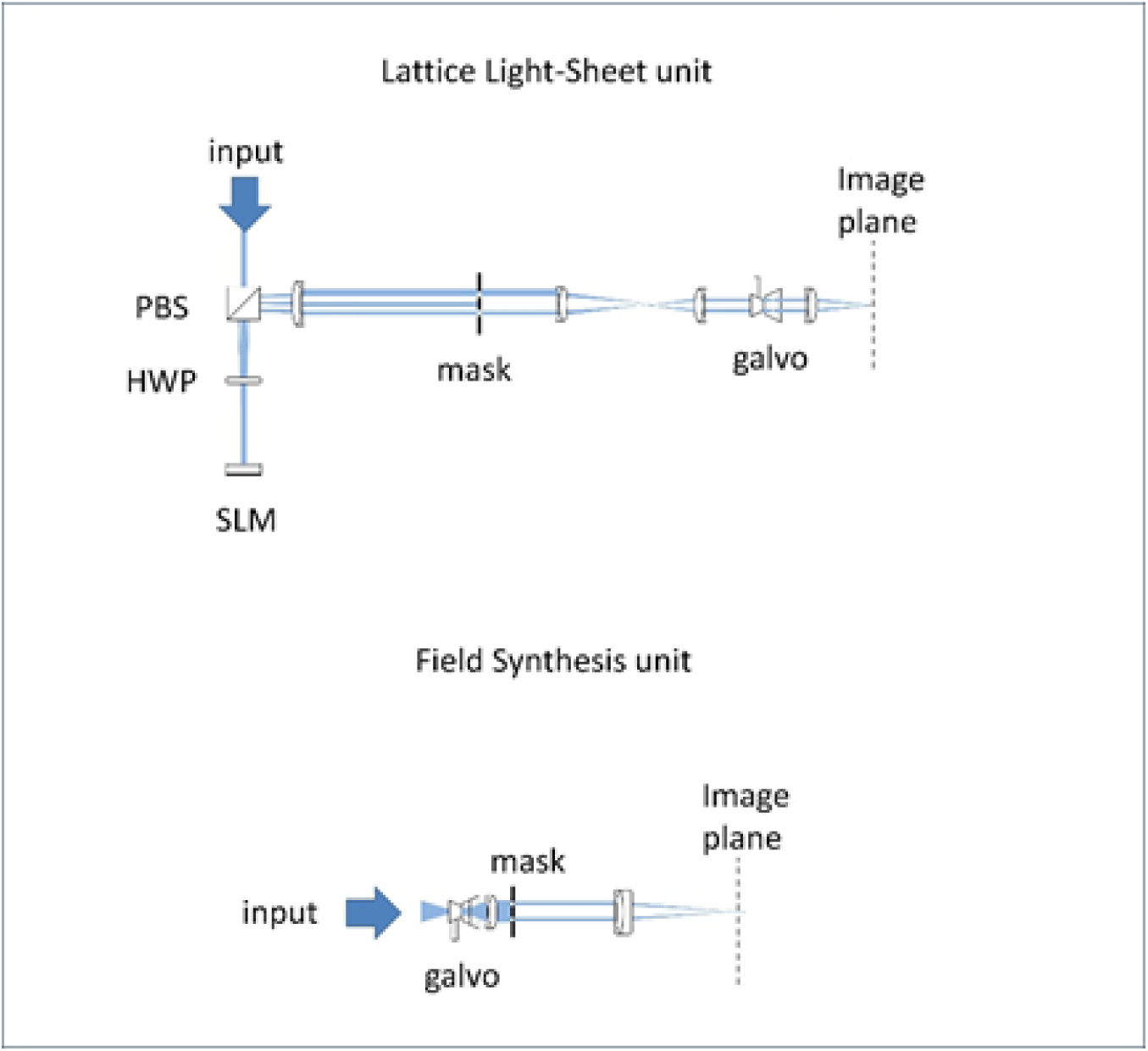
Basic components for conventional Lattice Light-Sheet and Field Synthesis Light-Sheet Generation. Units to produce lattice-light-sheets conventionally and with Field Synthesis. Both units take a line shaped laser beam as an input and output the final light-sheet into an image plane. PBS: Polarizing beam splitter, SLM: spatial light modulator; HWP: half wave plate.

**Figure S2.**
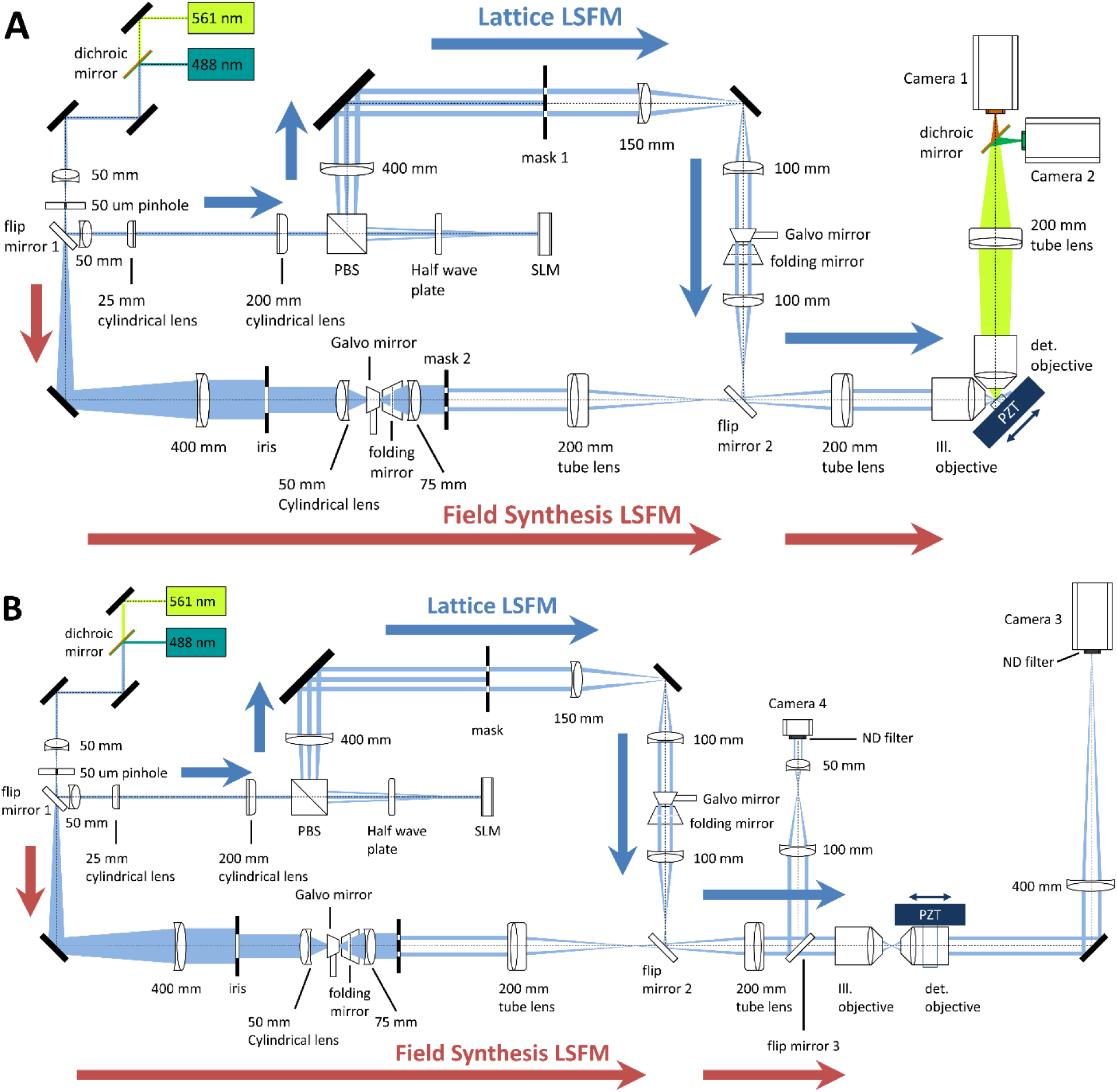
Schematic drawing of the experimental setup. (A) Microscope in the fluorescence imaging mode. (B) Microscope in the transmission imaging mode. The setup consists of two illumination paths, one (along the blue arrows) for conventional LLSM1 and the other (along the red arrows) for Field Synthesis LSFM. The optical path is selected with flip mirror 1 and 2. For routine inspection of the illumination wavefront, flip mirror 3 (not shown in A) redirects the light to camera 4, which is conjugate to the back-pupil plane of the excitation objective.

**Figure S3.**
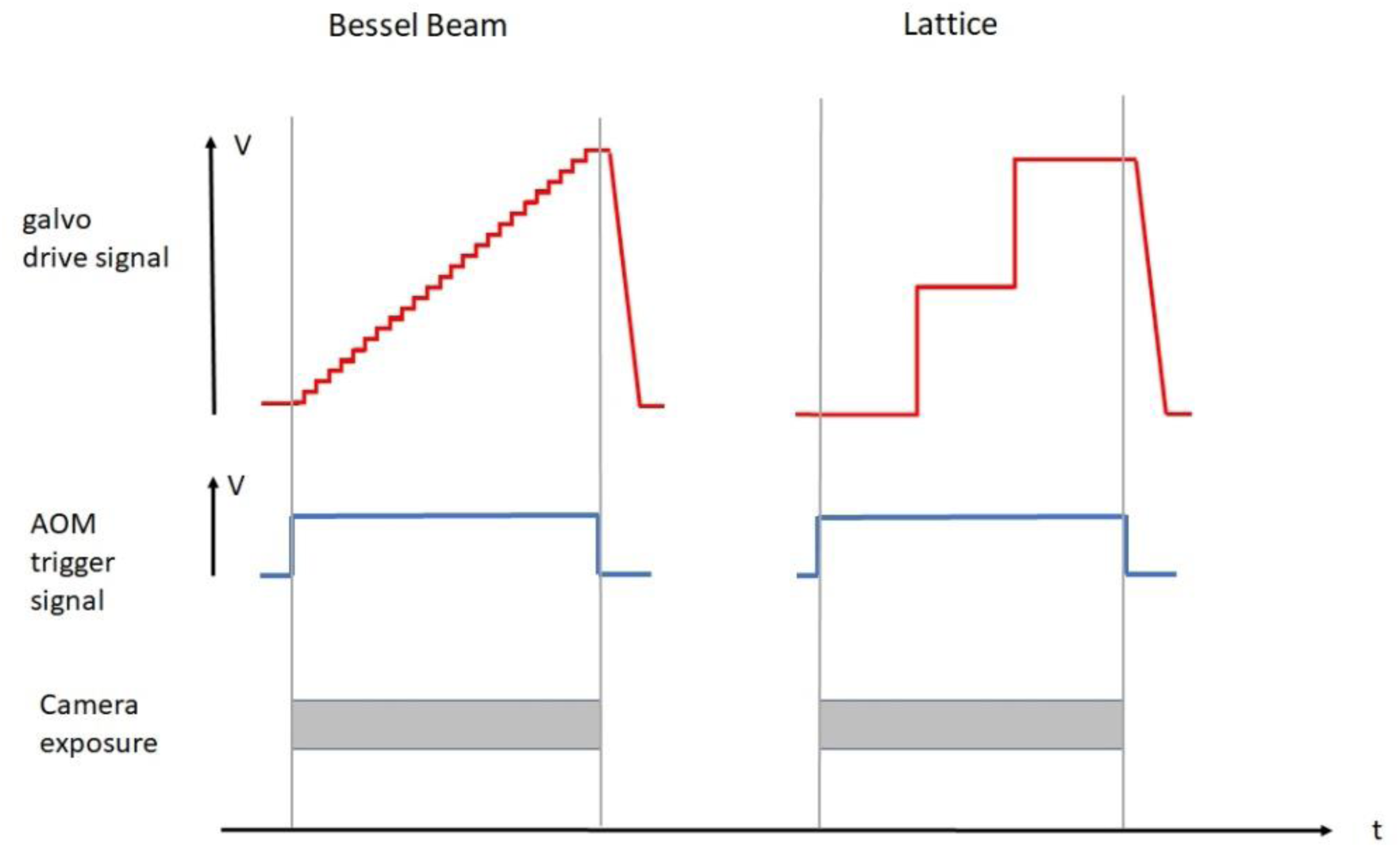
Timing diagram for Field synthesis. For Bessel beam light-sheets (or any light-sheet that has a continuous pupil function), the Galvo mirror in the Field Synthesis illumination train (see also Fig S1 and S2) is driven with a sawtooth signal with fine step size. For lattice light-sheets, the galvo is driven by a sawtooth signal that consists of two steps.

**Figure S4.**
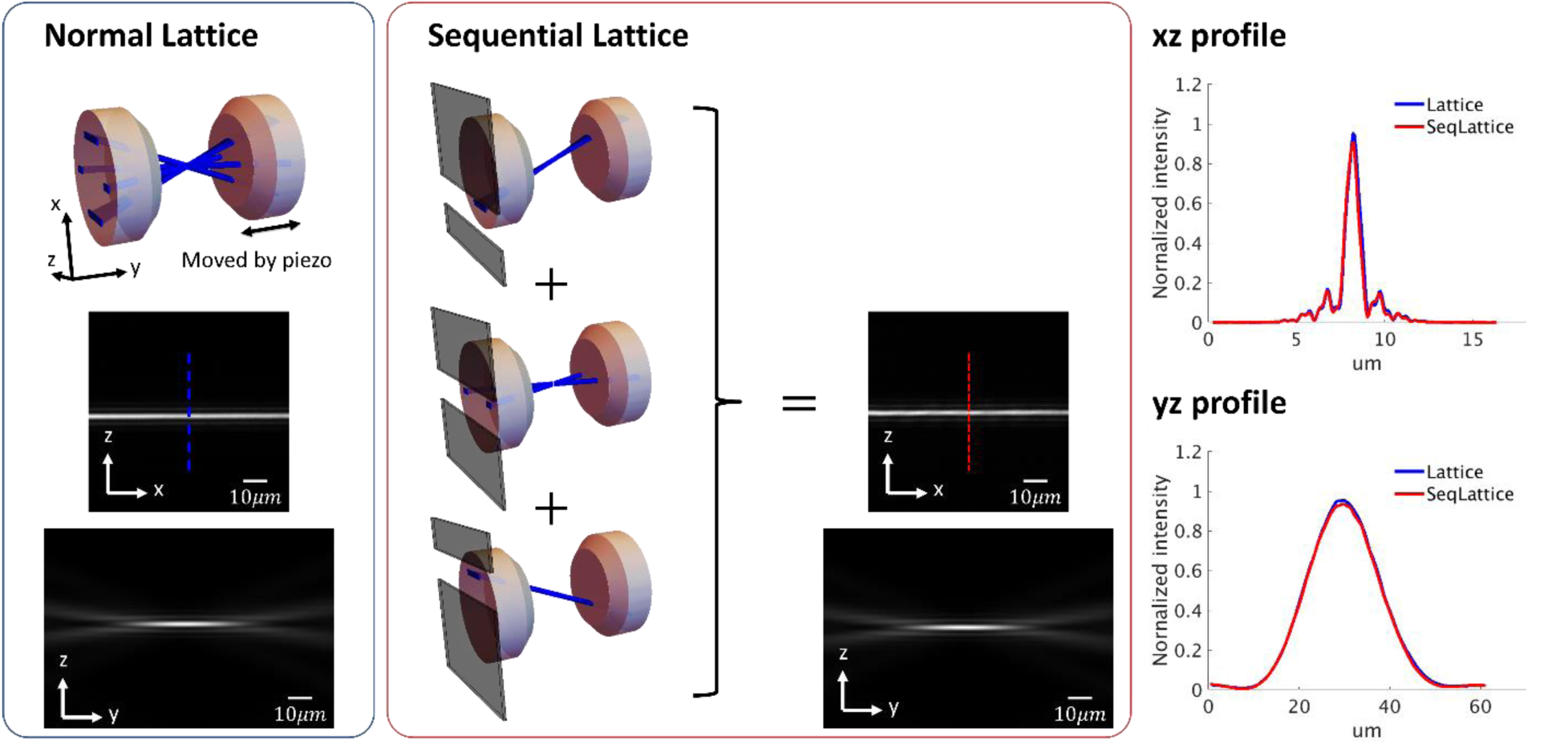
Sequential lattice experiment. This experiment enables testing the Field synthesis theorem using a conventional lattice light-sheet microscope. In the sequential lattice illumination mode, which simulates the Field Synthesis process, three diffraction orders of a square lattice pattern were individually selected by a slit mask on the back aperture (Fourier space) of the objective. Each order was separately imaged in image space and the three data sets were added. The xz image shows the light-sheet on the focal plane. The yz image shows the light-sheet along the propagation direction (y). The xz and yz profiles of the light-sheets are the averages of six line-profiles from xz and yz images. The cross-sectional profiles show a comparison between dithered, normal square lattice light-sheets and the results obtained from the sequential lattice approach.

**Figure S5.**
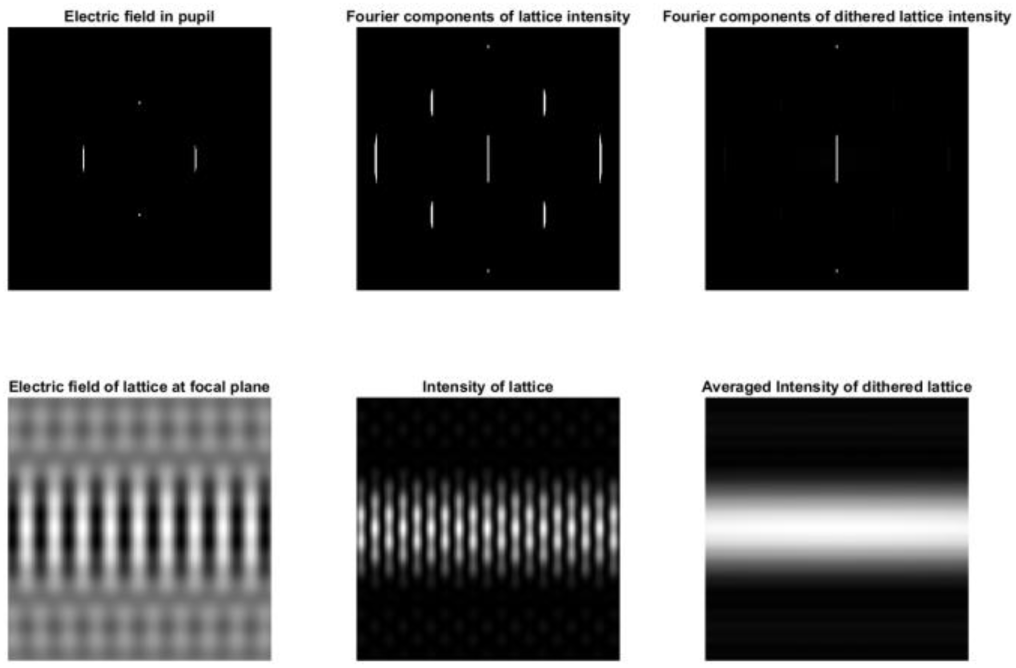
Simulation of the Electric field and the Intensity of a square lattice light-sheet in Fourier and real space. Top row from left to right: Electric field at the pupil created by the square lattice mask, Magnitude of the Fourier transform of the intensity of the square lattice, Magnitude of the Fourier transform of the intensity of the dithered square lattice. Bottom row from left to right: Electric field of the lattice at the focal plane, Intensity (square modulus) of the lattice, Intensity of the square lattice dithered over one period. The figure was generated by FieldSynthesisVersusLattice.m.

**Figure S6.**
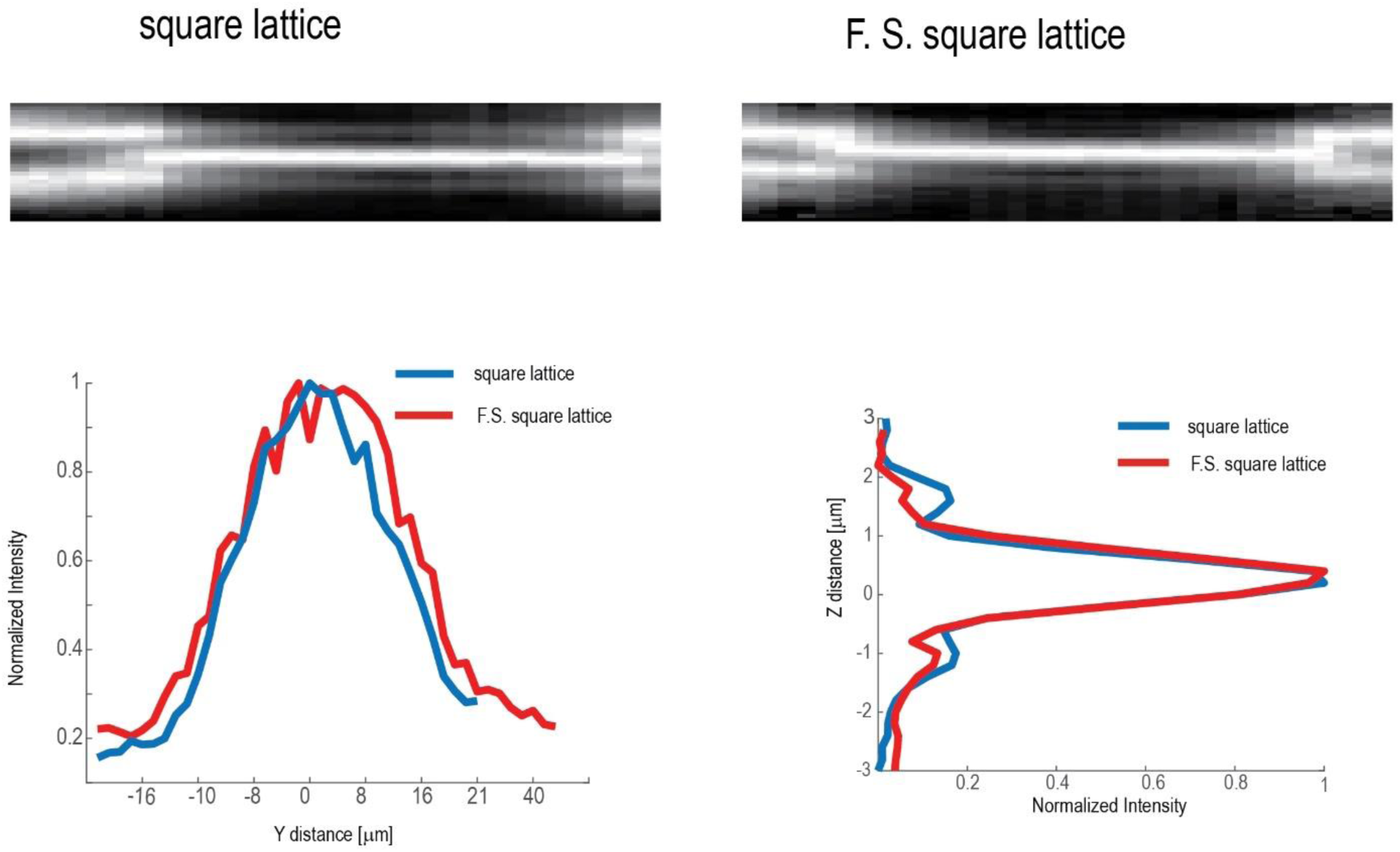
Measurement of the intensity distribution of a square lattice and Field Synthesis light-sheet in the sample plane. A 50nm fluorescent bead was scanned through the light-sheets and at every scan position, the measured fluorescence was summed up, resulting in the measurements shown in the top row. Bottom row shows intensity profiles along the propagation and axial direction of the light sheet.

**Figure S7.**
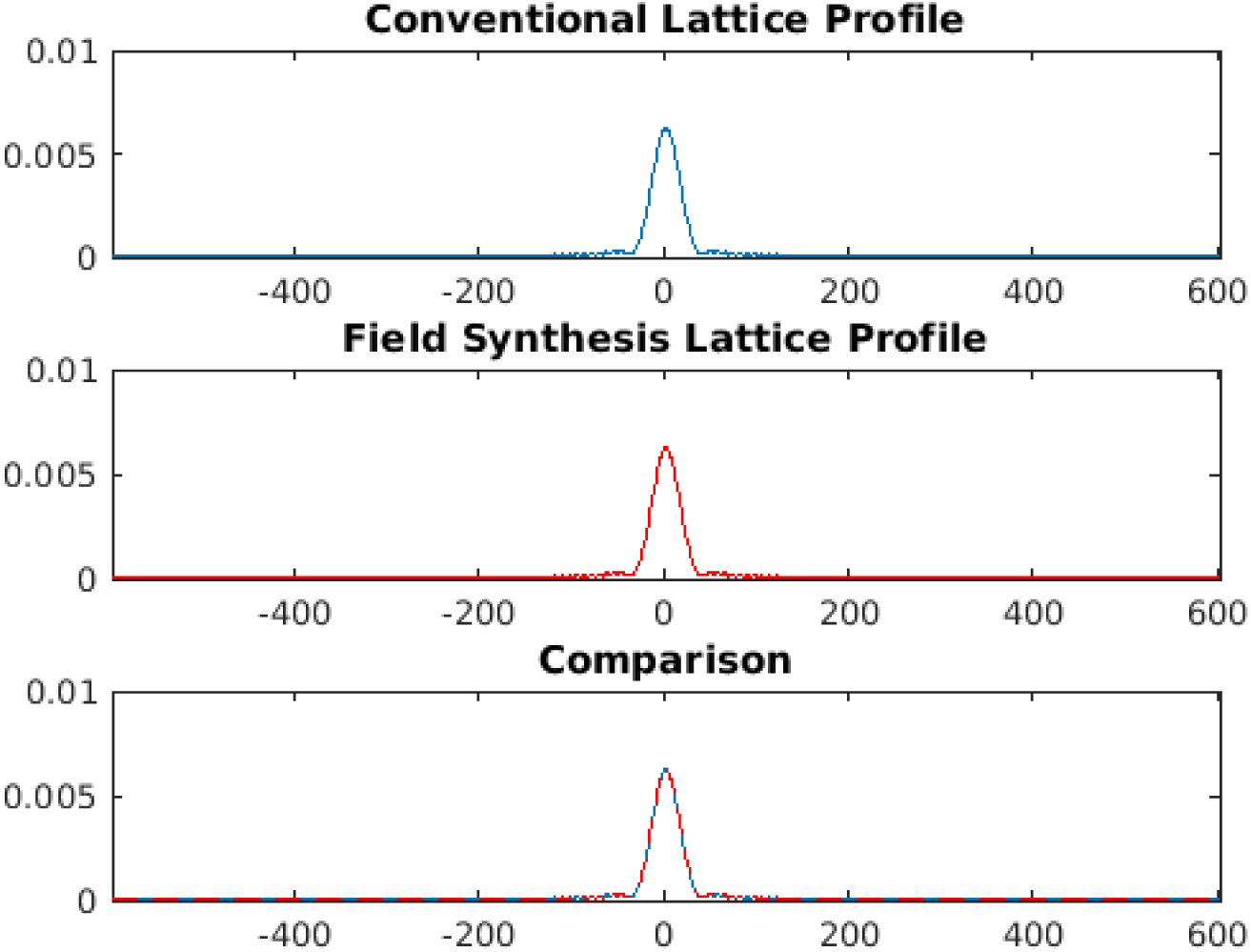
Simulation of conventional square Lattice profile and Field Synthesis square Lattice profile. Top: Conventional lattice profile generated by dithering the lattice intensity. Middle: Field synthesis lattice profile generated by summing line scans over the pupil mask. Bottom: Overlay of the conventional and field synthesis lattice profiles. The figure was generated by FieldSynthesisVersusLattice.m

**Figure S8.**
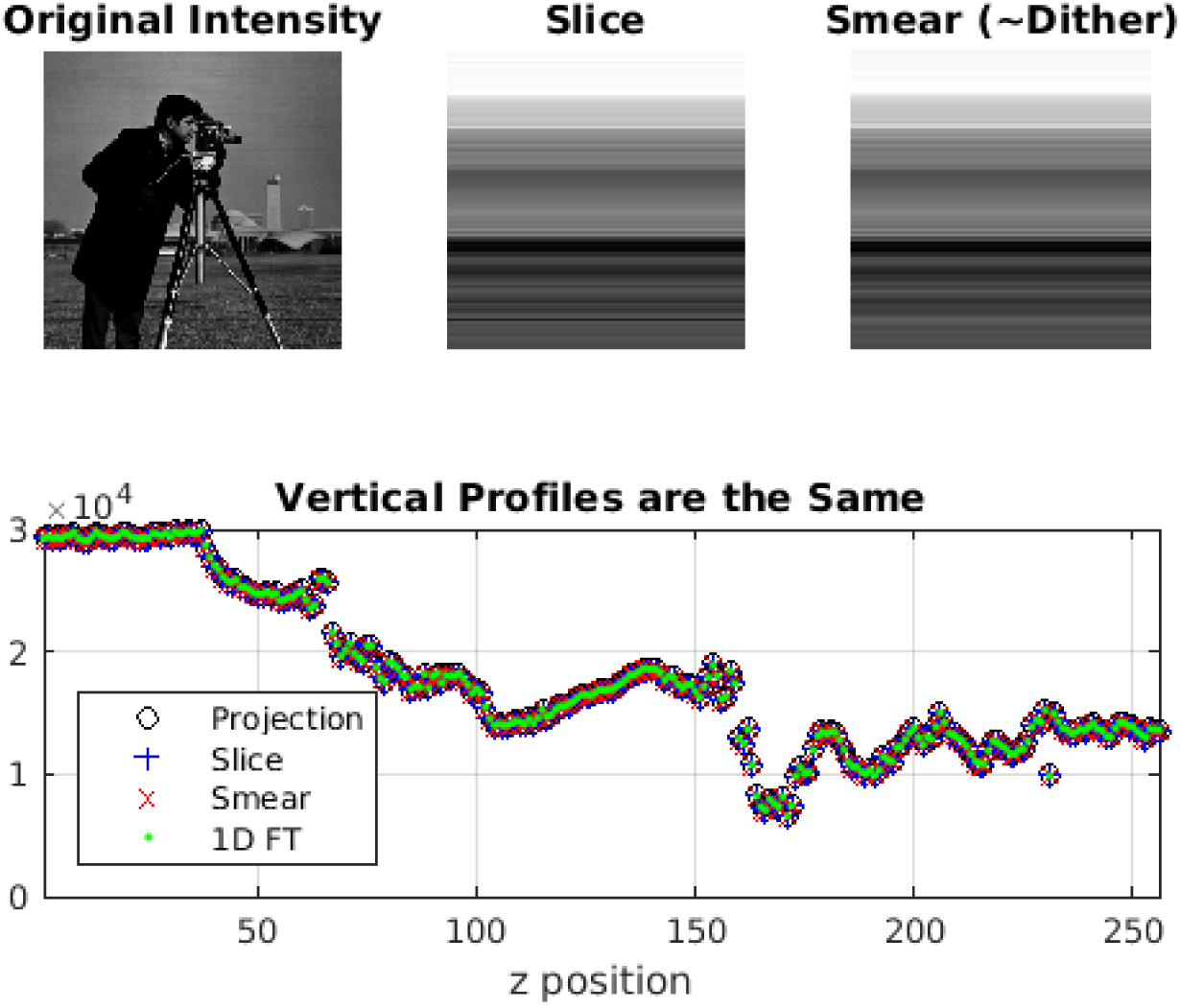
Simulation and Proof of Concept for Field Synthesis. Top from left to right: Original intensity pattern created by a complex mask, Sum of the square modulus of slices of the pupil mask as in the field synthesis method, Smearing of the intensity pattern as done in dithering. The bottom plot shows the overlay of several methods of calculating the vertical profile created by slicing or smearing. Projection is the profile created by averaging each row of the original intensity. Slice is the profile created by the field synthesis method. Smearing is the profile created by dithering the intensity. 1D FT is the profile created by taking a one-dimensional Fourier transform of the pupil mask in the z-direction and then summing the square modulus in the x-direction as in the above proof. Created by FieldSythesisTheorem.m.

**Figure S9.**
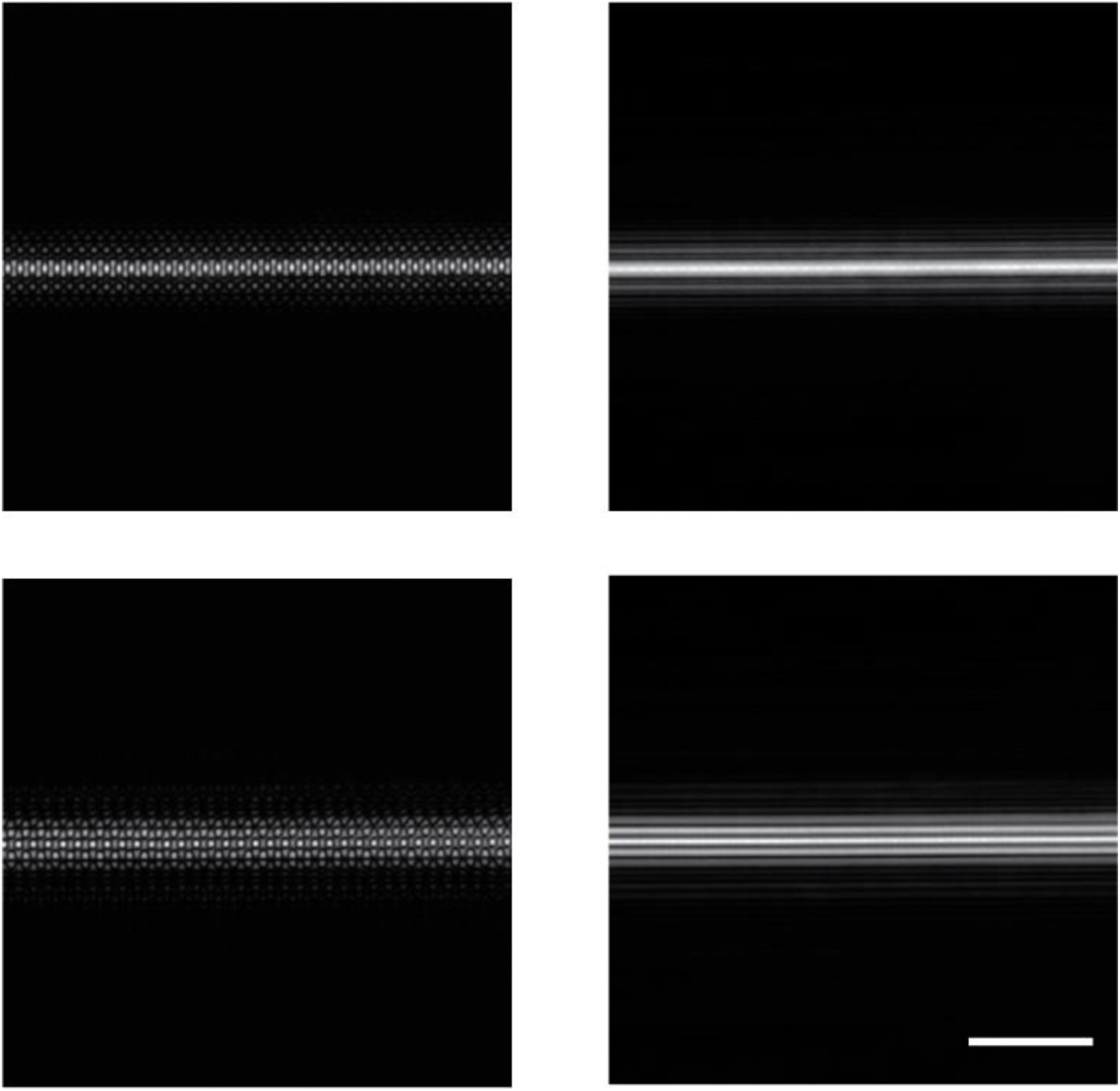
Square (top) and hexagonal (bottom) lattice light-sheets as measured in transmission produced by our experimental setup shown in Figure S2. Undithered lattices are shown in the left column and the dithered lattices in the right column. The undithered lattices are shown here as a quality control for our experimental setup. Typical interference patterns of a 2D optical lattice are clearly visible (see also Chen et al^1^). Scale bar: 10 microns

## Supplementary Tables

**Supporting Table 1.** Imaging parameters

| name | Camera exposure | Number of z-slices | Stack acquisition time | Laser power in pupil | NA<sub>id</sub> | NA<sub>od</sub> |
| --- | --- | --- | --- | --- | --- | --- |
| Scanned Bessel beam beads | 10ms | 101 | 1.011s | 3.84 mW | 0.439 | 0.545 |
| Field synthesis Bessel beam beads | 10ms | 101 | 1.011s | 1.47 mW | 0.444 | 0.533 |
| Scanned Bessel beam EB3 | 20 ms | 376 | 7.5s | 210μW | 0.439 | 0.545 |
| Field Synthesis Bessel beam EB3 | 20ms | 326 | 6.5s | 140μW | 0.444 | 0.533 |
| Square lattice MV3 | 10ms | 201 | 4.4s | 75μW @488nm<br>30μW @561nm | 0.439 | 0.545 |
| Field Synthesis MV3 | 10ms | 201 | 2.13 | 380μW @488nm<br>100μW @561 | 0.438 | 0.539 |

**Supporting Table 2.** light-Sheet parameters. Table lists mean and standard deviation as measured over six positions across each light-sheet dataset. Last column lists the total number of measurements (i.e. 18 equals to three light-sheet data sets with six measurement positions on each) used to compute average and standard deviation.

| Light-sheet type | Axial FWHM | Propagation FWHM | # for average |
| --- | --- | --- | --- |
| Scanned Bessel Beam | $0.34 \pm 0.015 \mu m$ | $13.27 \pm 0.119 \mu m$ | 18 |
| Field Synthesis Bessel beam | $0.34 \pm 0.017 \mu m$ | $13.16 \pm 0.085 \mu m$ | 18 |
| Squared lattice | $0.89 \pm 0.021 \mu m$ | $17.71 \pm 0.261 \mu m$ | 24 |
| Field Synthesis squared lattice | $0.92 \pm 0.010 \mu m$ | $16.80 \pm 0.403 \mu m$ | 24 |
| Hexagonal lattice | $0.37 \pm 0.003 \mu m$ | $10.89 \pm 0.210 \mu m$ | 18 |
| Field Synthesis hexagonal lattice | $0.38 \pm 0.015 \mu m$ | $10.90 \pm 0.739 \mu m$ | 24 |
| Sequential square lattice | $0.89 \pm 0.058 \mu m$ | $18.00 \pm 0.740 \mu m$ | 12 |

## Supplementary Movies

**Movie S1.**
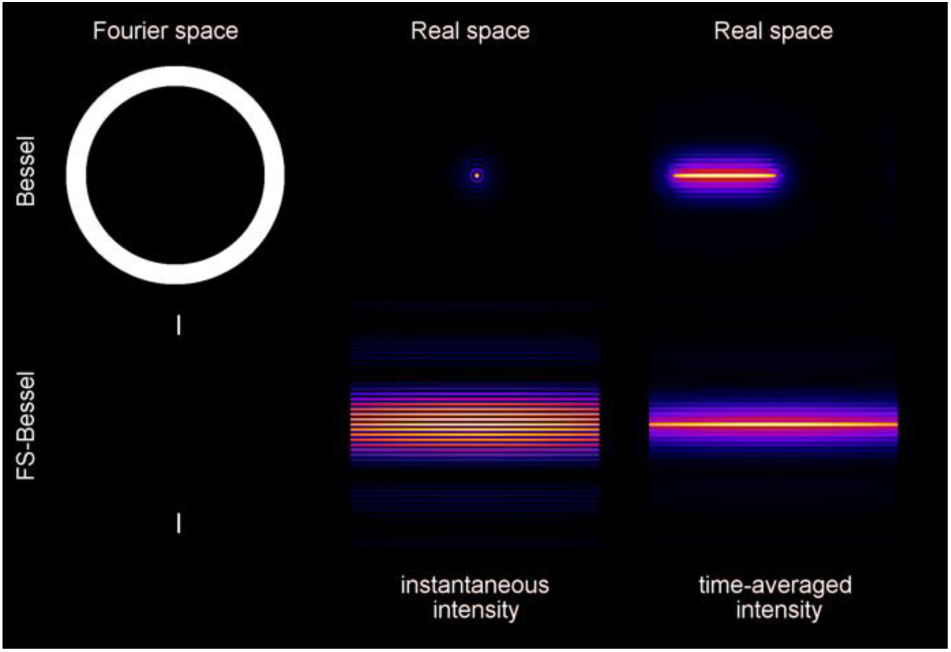
Simulation of light-sheet generation. The top row of the simulation shows an annular mask that is placed in a Fourier plane, which produces a Bessel beam in sample space in this example. In the middle column on top, the instantaneous intensity of a Bessel beam in real space is shown as it is scanned laterally. On the right, the time averaged intensity is shown. On the bottom, the Field Synthesis process is shown. On the left, a line scan over the pupil mask is shown and the resulting instantaneous interference pattern in real space is shown in the middle. On the right, the time averaged intensity is shown.

**Movie S2.**
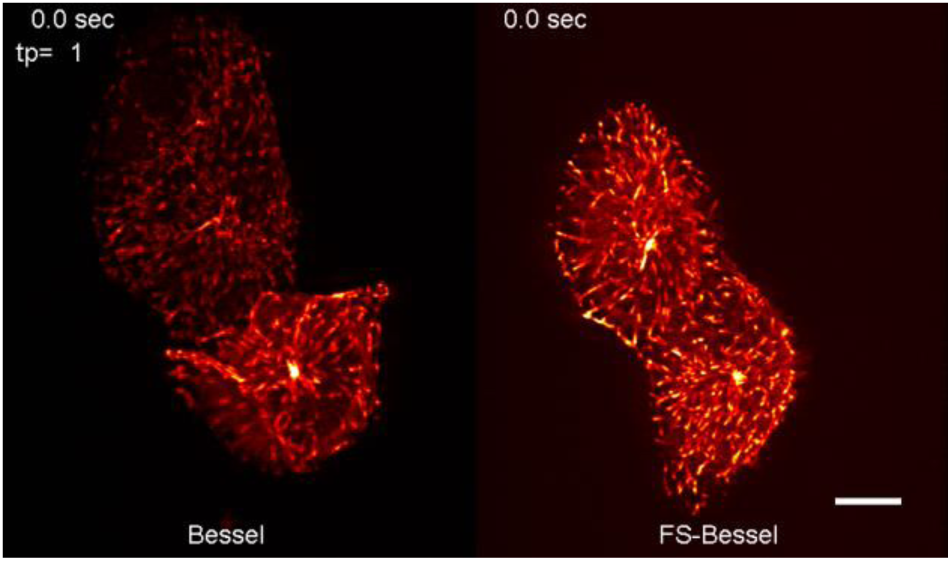
EB3 Dynamics Imaged with Bessel beam- and Field Synthesis Bessel beam-Light-sheet Microscopy. U2OS Cell was labeled with EB3-mNeonGreen. When imaged by scanned Bessel beam light-sheet microscopy (left), the camera exposure time was set to 20ms and the volumetric acquisition rate was 7.5s per volume. When imaged by Field Synthesis Bessel beam light-sheet microscopy (right), the camera exposure time was set to 20ms and the volumetric acquisition rate was 6.5s per volume. Scale bar 10 microns.

**Movie S3.**
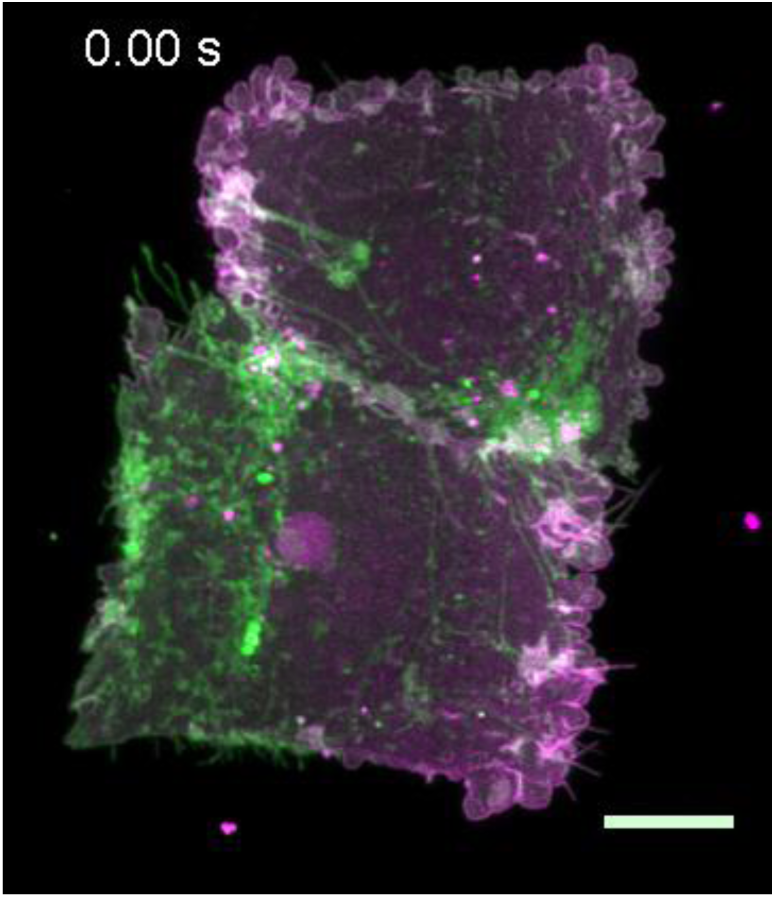
3D timelapse imaging of an MV3 cell imaged with square lattice light-sheet microscopy. An MV3 labeled with GFP AktPH biosensor and CAAX-Tdtomato membrane. Exposure time was 10ms per frame and total stack acquisition time amounted to 4.43s. Biosensor is shown in magenta and membrane in green. Scale bar 10 microns.

**Movie S4.**
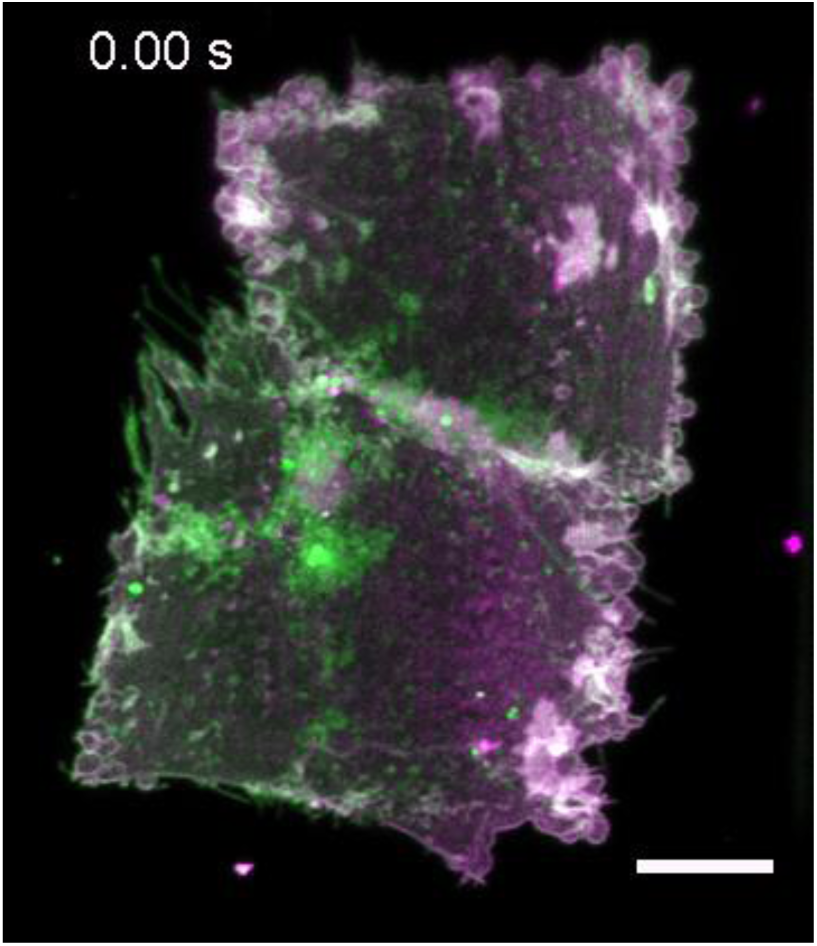
3D timelapse imaging of an MV3 cell imaged with Field Synthesis square lattice light-sheet microscopy. The same MV3 as shown in movie S3, imaged after the square lattice acquisition. Exposure time was 10ms per frame and total stack acquisition time amounted to 2.13s. Biosensor is shown in magenta and membrane in green. Scale bar 10 microns.

**Movie S5.**
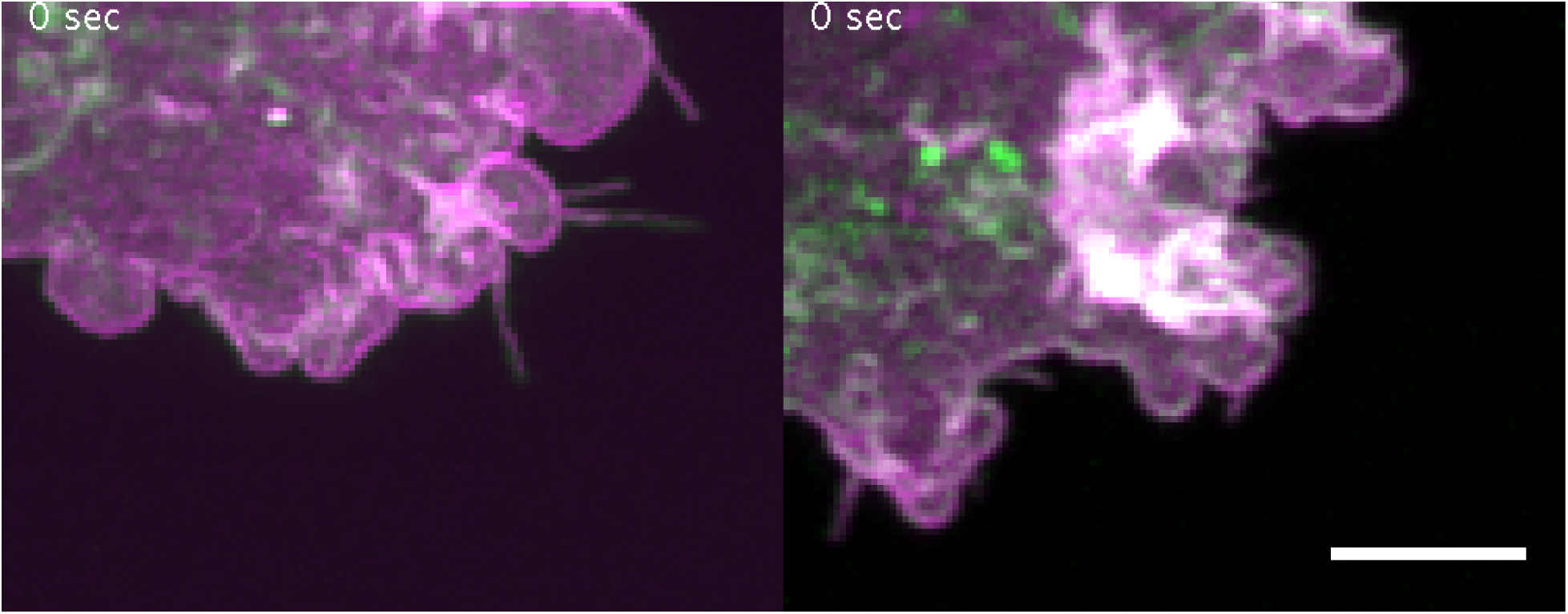
Bleb and filopodia dynamics imaged with square lattice and Field Synthesis square lattice light-sheet microscopy. The same MV3 cell as shown in movie S4 and S5, but a zoomed in view on the bottom right corner of the cell. Left side shows 14 timepoints acquired with square lattice and right side shows 28 time points acquired with Field Synthesis, spanning ~57s. Scale bar 5 microns.

## References

1. Davidovits, P. & Egger, M.D. Scanning Laser Microscope. Nature 223, 831–831 (1969).

2. Laissue, P.P. Alghamdi, R.A., Tomancak, P., Reynaud, E.G. & Shroff, H. Assessing phototoxicity in live fluorescence imaging. Nature Methods 14, 657–661 (2017).

3. Cranfill, P.J. et al. Quantitative assessment of fluorescent proteins. Nature Methods 13, 557–562 (2016).

4. Stelzer, E.H.K. Light-sheet fluorescence microscopy for quantitative biology. Nature Methods 12, 23–26 (2015).

5. Chen, B.-C. et al. Lattice light-sheet microscopy: Imaging molecules to embryos at high spatiotemporal resolution. Science 346 (2014).

6. Hanser, B.M. Gustafsson, M.G., Agard, D.A. & Sedat, J.W. Phase-retrieved pupil functions in wide-field fluorescence microscopy. J Microsc 216, 32–48 (2004).

7. Gao, L. et al. Noninvasive Imaging beyond the Diffraction Limit of 3D Dynamics in Thickly Fluorescent Specimens. Cell 151, 1370–1385 (2012).

8. Planchon, T.A. et al. Rapid three-dimensional isotropic imaging of living cells using Bessel beam plane illumination. Nature Methods 8, 417–423 (2011).

9. Vettenburg, T. et al. Light-sheet microscopy using an Airy beam. Nature Methods 11, 541–544 (2014).

10. Fahrbach, F.O. Gurchenkov, V., Alessandri, K., Nassoy, P. & Rohrbach, A. Self-reconstructing sectioned Bessel beams offer submicron optical sectioning for large fields of view in light-sheet microscopy. Optics Express 21 (2013).

11. Quirin, S. et al. Calcium imaging of neural circuits with extended depth-of-field light-sheet microscopy. Optics Letters 41 (2016).

12. Gao, L. Extend the field of view of selective plan illumination microscopy by tiling the excitation light sheet. Optics Express 23 (2015).

13. Valm, A.M. et al. Applying systems-level spectral imaging and analysis to reveal the organelle interactome. Nature 546, 162–167 (2017).

## Supplemental References

1. Chen, B.-C. et al. Lattice light-sheet microscopy: Imaging molecules to embryos at high spatiotemporal resolution. Science 346, 1257998–1257998 (2014).

2. Kner, P., Chhun, B. B., Griffis, E. R., Winoto, L. & Gustafsson, M. G. L. Super-resolution video microscopy of live cells by structured illumination. Nat. Methods 6, 339–42 (2009).

3. Schindelin, J. et al. Fiji: an open-source platform for biological-image analysis. Nat. Methods 9, 676–682 (2012).

4. Kirshner, H., Aguet, F., Sage, D. & Unser, M. 3-D PSF fitting for fluorescence microscopy: implementation and localization application. J. Microsc. 249, 13–25 (2013).

5. Dean, K. M. et al. Imaging subcellular dynamics with fast and light-efficient volumetrically parallelized microscopy. Optica 4, 263 (2017).

6. Dean, K. M. M. et al. Diagonally Scanned Light-Sheet Microscopy for Fast Volumetric Imaging of Adherent Cells. Biophys. J. 110, 1456–65 (2016).

7. Lanni, F. et al. Excitation field synthesis as a means for obtaining enhanced axial resolution in fluorescence microscopes Bioimaging 1.4 187–196 (1993).

